# Characterization of a haplotype-reference panel for genotyping by low-pass sequencing in Swiss Large White pigs

**DOI:** 10.1101/2021.03.11.434987

**Authors:** Adéla Nosková, Meenu Bhati, Naveen Kumar Kadri, Danang Crysnanto, Stefan Neuenschwander, Andreas Hofer, Hubert Pausch

## Abstract

**Background:** The key-ancestor approach has been frequently applied to prioritize individuals for whole-genome sequencing based on their marginal genetic contribution to current populations. Using this approach, we selected 70 key ancestors from two lines of the Swiss Large White breed that have been selected divergently for fertility and fattening traits and sequenced their genomes with short paired-end reads.

**Results:** Using pedigree records, we estimated the effective population size of the dam and sire line to 72 and 44, respectively. In order to assess sequence variation in both lines, we sequenced the genomes of 70 boars at an average coverage of 16.69-fold. The boars explained 87.95 and 95.35% of the genetic diversity of the breeding populations of the dam and sire line, respectively. Reference-guided variant discovery using the GATK revealed 26,862,369 polymorphic sites. Principal component, admixture and F_ST_ analyses indicated considerable genetic differentiation between the lines. Genomic inbreeding quantified using runs of homozygosity was higher in the sire than dam line (0.28 vs 0.26). Using two complementary approaches (CLR and iHS), we detected 51 signatures of selection. However, only six signatures of selection overlapped between both lines. We used the sequenced haplotypes of the 70 key ancestors as a reference panel to call 22,618,811 genotypes in 175 pigs that had been sequenced at very low coverage (1.11-fold) using GLIMPSE. The genotype concordance, non-reference sensitivity and non-reference discrepancy between thus inferred and Illumina PorcineSNP60 BeadChip-called genotypes was 97.60, 98.73 and 3.24%, respectively. The low-pass sequencing-derived genomic relationship coefficients were highly correlated (r > 0.99) with those obtained from microarray genotyping.

**Conclusions:** We assessed genetic diversity within and between two lines of the Swiss Large White pig breed. Our analyses revealed considerable differentiation, even though the split into two populations occurred only few generations ago. The sequenced haplotypes of the key ancestor animals enabled us to implement genotyping by low-pass sequencing which offers an intriguing cost-effective approach to increase the variant density over current array-based genotyping by more than 350-fold.

## Background

Swine production follows the classical breeding pyramid. Genetic gain is generated in nucleus herds and transmitted via the multiplier to the production unit. Swiss pig production relies on maternal and paternal Swiss Large White (SLW) lines at the top level of the breeding pyramid. For decades, the SLW breed has been maintained as a universal breed, selected for production and fertility traits. In 2002, the population was divided into sire and dam lines that have been divergently selected for fattening and reproduction since then. Approximately 32.5% and 30% of the genes of 2.5 million fattening pigs slaughtered in 2020 in Switzerland originate from the dam and sire line, respectively. Both lines are maintained in purebred nucleus herds. However, little is known about the genetic diversity within the lines.

The SLW breeding boars are selected based on genome-based breeding values that are predicted using genotypes obtained with a customized version of the Illumina PorcineSNP60 BeadChip. Apart from a small number of putatively causal variants that are included in the custom part, the content of the currently used microarray was designed in a way that it is useful for mainstream breeds [1]. However, the genetic constitution of the SLW breed beyond the microarray-derived SNP remains largely unknown. The sequencing of key ancestor animals has been proposed as a cost-efficient way to assess sequence variation within a population [2]. Due to the use of individual boars in artificial insemination and intense selection in nucleus herds, the effective population size of most pig breeding populations is low. Thus, most common polymorphic sites segregating in the population can be traced back to the genomes of important contributors to the current population [3, 4]. The key ancestor approach was frequently applied to identify the most important contributors to current cattle breeding populations [4]. Recently it was also used to prioritize animals for sequencing in commercial pig breeding lines [5].

The availability of sequence variant genotypes from key ancestor animals enables imputing sequence-level genotypes for animals that had been genotyped at lower density [6–8]. In livestock populations that are routinely genotyped using 60K genotyping arrays, sequence variant genotypes are typically imputed using stepwise imputation [9]. In a first step, 60K genotypes are imputed to higher density (e.g., 700K) using animals that have been genotyped with high-density genotyping arrays. In a second step, the partially imputed high-density genotypes are imputed to the sequence level based on a sequenced reference panel. The accuracy of imputing 60K genotypes directly to the sequence level is low, particularly for rare variants, rendering most of them uninformative for downstream analyses such as genomic prediction and association testing [10, 11]. Reference-guided variant phasing and imputation from low-pass sequencing data offers an intriguing alternative approach to the two-step imputation approach in pedigreed populations [12]. This approach utilises a sequenced haplotype reference panel that represents the diversity of the target population. Sequence variant genotypes of animals sequenced at very shallow coverage are then inferred conditional on the observed haplotypes of the reference panel. This method is particularly useful in species for which dense microarray-derived genotypes are not available. Recent investigations [6, 13, 14] suggest that a sequencing coverage less than 1-fold is sufficient to accurately infer genotypes at known loci - provided an informative haplotype reference panel is available.

Here we obtain whole-genome sequencing data from key ancestor animals to characterize genetic diversity, population structure, and signatures of selection in two divergently selected commercial pig breeds. Using the haplotypes of the key ancestor animals as a reference panel, we accurately genotype more than 22 million variants in animals that have been sequenced at low coverage.

## Results

Using pedigree-derived inbreeding coefficients, we estimated the effective population size of the sire and dam line of the SLW breed to 44 and 72, respectively. In order to assess sequence variation within the two lines, we prioritized 70 boars for whole-genome sequencing based on their marginal genetic contributions to the active breeding populations with a key ancestor approach. Of the 70 boars, 38 and 32 represent the sire and dam line, respectively, explaining 95.35 and 87.95% of the genetic diversity of the active breeding populations.

Following quality control (removal of adapter sequences, reads and bases of low sequencing quality), between 81.15 and 377.01 million read pairs (2 × 150 bp) per sample (mean: 165.55 ± 60.32 million read pairs) were aligned to the SSC11.1 assembly of the porcine genome. Using reads with high mapping quality (reads with mapping quality < 10 and SAM bitwise flag 1796 were not considered), the average sequencing coverage of the 70 boars was 16.69 ± 5.93-fold across all autosomes. Raw sequence read data of 70 pigs have been deposited at the European Nucleotide Archive (ENA) of the EMBL at BioProject PRJEB38156 and PRJEB39374.

A reference-guided multi-sample variant discovery and genotyping approach yielded genotypes at 28,407,060 sites (22,191,375 biallelic SNP, 4,379,470 biallelic INDEL, and 1,836,215 others, Table 1). We applied GATK’s VariantFiltration module for site-level hard filtration using parameters recommended in the best practice guidelines [15]. Subsequently, we applied Beagle (version 4.1; [16]) phasing and imputation to improve the genotype calls from GATK and to impute sporadically missing genotypes. Following the imputation, we retained 26,862,369 variants including 21,592,583 SNP and 5,269,786 INDEL. The number of polymorphic sites that were seen in the heterozygous (singletons) and homozygous (doubletons) state only once was 2,026,088 (7.54%) and 72,100 (0.27%), respectively. To prevent bias resulting from flawed genotypes in repetitive regions, we excluded 1,710,337 variants for which an excess of sequencing coverage was evident for downstream analyses. The transition/transversion (Ti/Tv)-ratio estimated from filtered and imputed variants was 2.28.

**Table 1:**
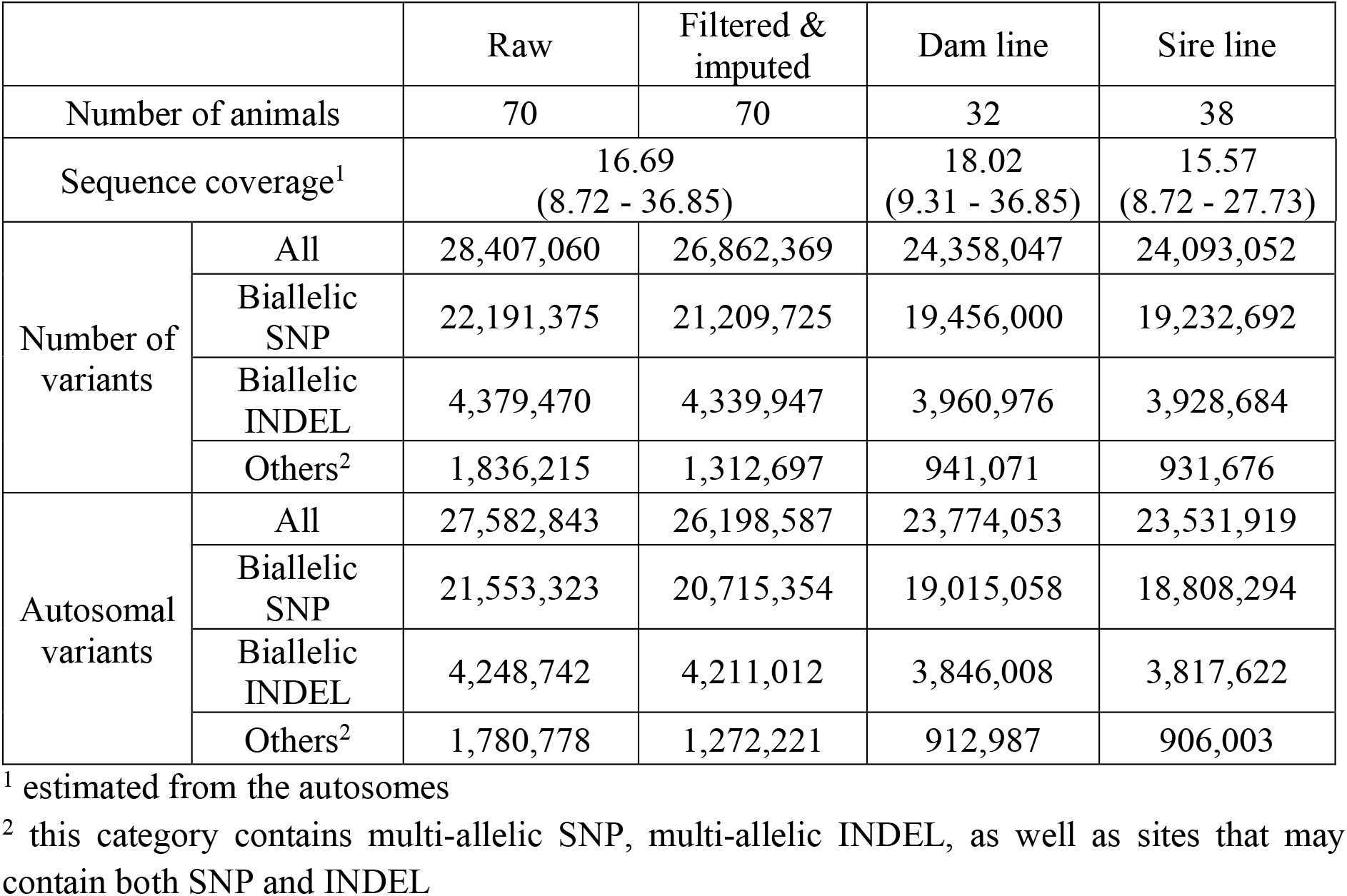
Variants detected in 70 sequenced key ancestor animals.

The resulting data were separated into two datasets containing 23,774,053 and 23,531,919 autosomal variants detected in 32 and 38 boars from the dam and sire line, respectively. Of the variants, 1,049,689 and 1,594,775 were fixed for the alternate allele in the dam and sire line, respectively. On average, we detected 11,119,760 ± 176,113 biallelic variants per animal (Figure 1A), of which 6,258,456 ± 280,127 and 4,861,304 ± 135,524 were heterozygous and homozygous for the reference allele, respectively. The average nucleotide diversity (π) across 452,444 overlapping windows (10 kb in size with 5 kb steps), spanning 22,840,217 and 22,529,446 biallelic variants, respectively, was 2.81 × 10^−3^ in the dam and 2.72 × 10^−3^ in the sire line.

**Figure 1.**
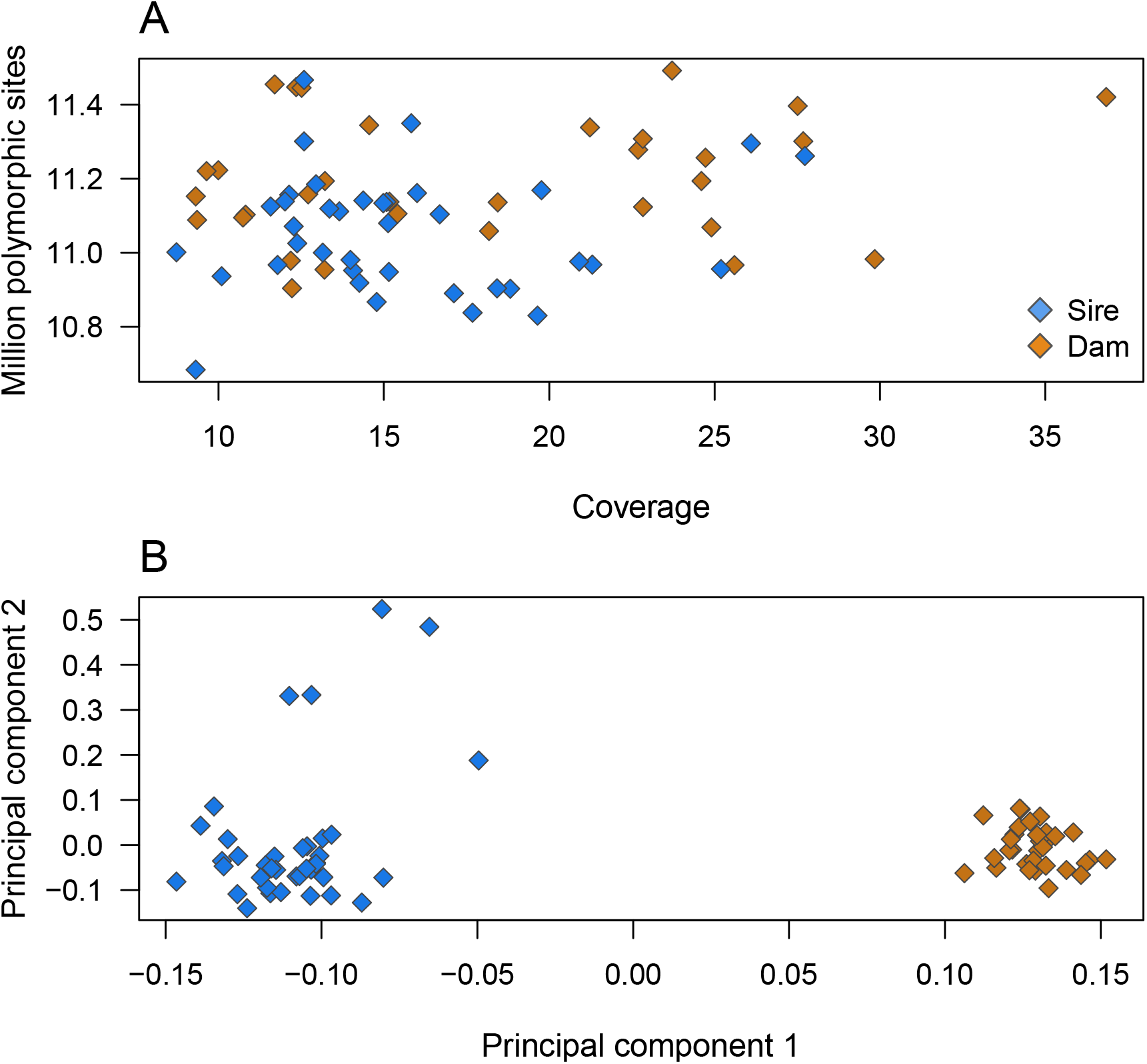
Sequencing of key ancestor animals from two pig lines. **(A)** Number of polymorphic sites detected in the 70 boars as a function of depth of coverage. **(B)** Plot of the first two principal components showing the separation of animals by breed and the relationship between both lines. Blue and orange symbols indicate 38 and 32 boars from the sire and dam line, respectively.

### Comparison between array-called and sequence-called genotypes

Sixty-eight boars (32 and 36 from the dam and sire line, respectively) that had average sequencing coverage between 8.72 and 36.85-fold (average: 16.79-fold) also had Illumina PorcineSNP60 BeadChip-called genotypes. Using the array-called genotypes at 54,600 autosomal SNP for which we were able to determine reference and alternate alleles as a truth set, we calculated genotype concordance, non-reference sensitivity and non-reference discrepancy between array-called and sequence-called genotypes as proposed by DePristo et al. [17]. Of the 54,600 SNP, 6,376 and 1,029 were fixed for the reference and alternate allele, respectively, and 47,195 were polymorphic in the array-called genotypes of the 68 pigs.

Of the 48,224 SNP that were either polymorphic or fixed for the alternate allele in the array-called genotypes, 46,009 (95.41%) and 45,951 (95.29%) were also present in the raw and filtered sequence variants, respectively. 1,232 SNP of the Illumina PorcineSNP60 BeadChip complement were missing in the sequenced set because they were either genotyped as INDEL or multiallelic sites using GATK and thus excluded from the comparison due to incompatible alleles. 983 and 1,041 SNP were not among the raw and filtered sequence variants, respectively, although the frequency of the minor allele was > 5% in the array-called genotypes for most (> 80%) of them. It is likely that these variants could not be matched with the sequence set due to either incompatible or ambiguous map coordinates.

Non-reference sensitivity was greater than 99% and non-reference discrepancy around 1% for the raw genotypes called by the GATK, suggesting that the high sequencing coverage facilitated accurate variant discovery (Table 2). The concordance between sequence-and array-called genotypes improved slightly after applying site-level hard filtration. Beagle phasing and imputation further increased the concordance and non-reference sensitivity as well as decreased the non-reference discrepancy of the filtered sequence variant genotypes.

**Table 2:**
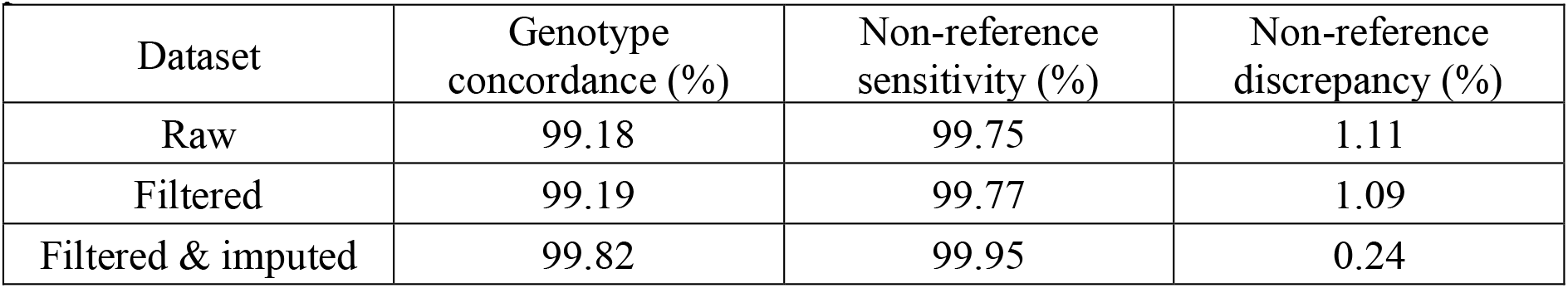
Comparison between sequence- and array-called genotypes at corresponding positions.

| Dataset | Genotype concordance (%) | Non-reference sensitivity (%) | Non-reference discrepancy (%) |
| --- | --- | --- | --- |
| Raw | 99.18 | 99.75 | 1.11 |
| Filtered | 99.19 | 99.77 | 1.09 |
| Filtered & imputed | 99.82 | 99.95 | 0.24 |

### Population structure and genetic diversity

To investigate the population structure, ancestry and genetic diversity among the 70 sequenced pigs, we performed principal component, admixture and F_ST_ analyses. The principal components were extracted from a genomic relationship matrix constructed from 23,691,198 autosomal sequence variants that had minor allele frequency greater than 0.01.

The first principal component of the genomic relationship matrix explained 8.61% of the variation and separated the animals by lines (Figure 1B). The second principal component explaining 2.68% of the variation revealed variability within the sire line. Five outlier animals along the second axis of variation descended from imported Large White boars, thus reflecting an introgression of foreign genes into the sire line.

We performed an admixture analysis using 1,207,189 independent biallelic SNP to assess gene flow between both lines. As expected, K = 2 was the most plausible number of genetically distinct clusters (Supplementary Fig. S1, Additional File 1). The cross-validation error for K = 1, K = 2 and K = 3 was 0.561, 0.546 and 0.564, respectively. At K = 2, six individuals (one from the dam and five from the sire line) showed some admixture.

In order to investigate if pronounced allele frequency differences exist between both lines, we performed a SNP-based genetic differentiation analysis using the Weir and Cockerham weighted F_ST_ statistic [18]. We observed multiple 10 kb sliding windows scattered throughout the genome with F_ST_ values greater than 0.25, indicating genetic divergence of both lines (Supplementary Fig. S2, Additional File 2). The average weighted F_ST_ value across all windows was 0.07.

We estimated runs of homozygosity (ROH) with BCFtools/ROH [19] based on GATK-derived Phred-scaled likelihoods for 19,146,365 biallelic SNP to investigate genomic inbreeding in both lines. In total 111,201 ROH with an average length of 391.28 kb (ranging from 50 kb to 11.1 Mb) were detected (Phred-scaled likelihood > 70). The ROH contained an average number of 3,176 SNP (ranging from 29 to 87,699). The boars from the dam and sire line had 1,604 ± 133 and 1,575 ± 91 ROH with an average size of 377,928 and 402,731 bp, respectively. The genomic inbreeding (F_ROH_, i.e., the fraction of the autosomal genome covered by ROH), was 0.26 ± 0.03 and 0.28 ± 0.03 for the dam and sire line, respectively. We classified the ROH into short (50 - 100 kb), medium (100 kb - 2 Mb) and long ROH (above 2 Mb) (Figure 2). Most ROH belonged to the medium length class. The average F_ROH_ was similar in both lines for small and medium ROH. However, F_ROH_ was higher for long ROH in the sire line.

**Figure 2.**
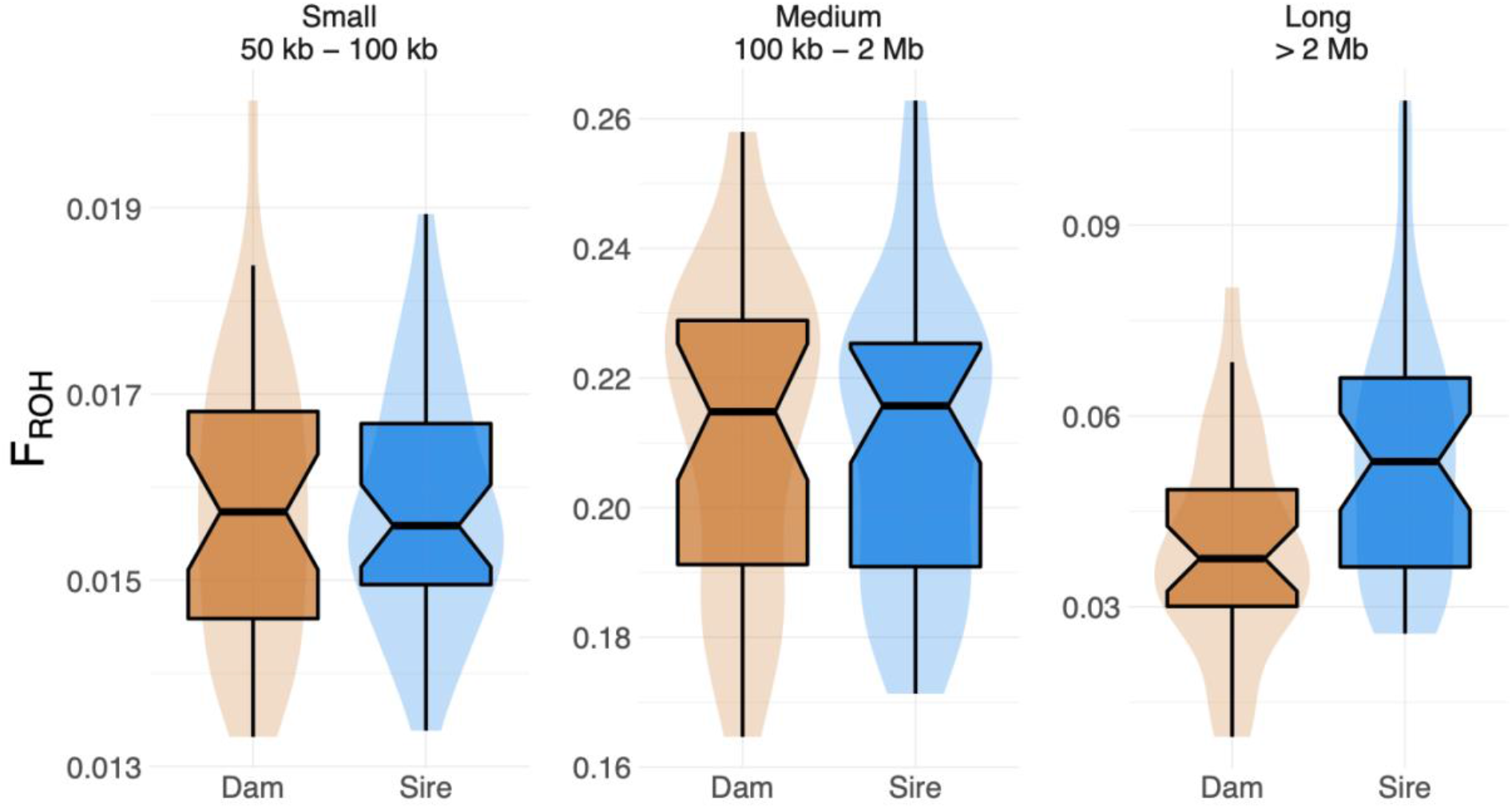
Genomic inbreeding in the two lines. F_ROH_ in dam and sire line, estimated for three groups of ROH classified based on their length: small (50 kb – 100 kb), medium (100 kb – 2 Mb) and long (> 2 Mb).

### Variant annotation

In 32 boars from the dam line, we annotated 23,774,053 (19,087,807 SNP; 4,038,170 INDEL) variants, including 2,567,754 variants that were not detected in the sire line. In 38 boars of the sire line, we annotated 23,531,919 (18,881,067 SNP; 4,009,043 INDEL) variants, including 2,325,620 that were not detected in the dam line. When compared to 63,832,658 germline variants listed for *Sus scrofa* in the Ensembl database (release 101), 5,745,790 (24.17%, dam line) and 5,693,068 (24.19%, sire line) variants were novel, of which the majority were INDEL and 14.66% and 14.64% were biallelic SNP.

We used the Ensembl Variant Effect Predictor software (VEP, release 98; [20]) to predict functional consequences for the sequence variants (Table 3). In total, 2.96% (dam line) and 2.94% (sire line) of the variants were in exons. Putative impacts of missense variants on protein function were predicted using the SIFT (sorting intolerant from tolerant) scoring algorithm [21] as implemented in the VEP software. The scoring algorithm classified 12,024 and 11,958 amino acid substitutions in the dam and sire line, respectively, as “deleterious” (SIFT score < 0.05).

**Table 3:** Predicted consequences of variants segregating in two lines. The table shows only the most sever consequence for a variant.

| Consequence type (most severe) | Dam line | Sire line |
| --- | --- | --- |
| Splice donor variant | 1,396 | 1,421 |
| Splice acceptor variant | 1,126 | 1,096 |
| Stop gained | 1,615 | 1,604 |
| Frameshift variant | 10,912 | 11,043 |
| Stop lost | 595 | 587 |
| Start lost | 423 | 421 |
| Inframe insertion | 990 | 987 |
| Inframe deletion | 1,164 | 1,186 |
| Protein altering variant | 62 | 62 |
| Missense variant | 70,758 | 69,983 |
| Splice region variant | 22,493 | 22,148 |
| Incomplete terminal codon variant | 12 | 11 |
| Synonymous variant | 76,977 | 75,279 |
| Stop retained variant | 149 | 135 |
| Start retained variant | 4 | 4 |
| Coding sequence variant | 98 | 96 |
| Mature miRNA variant | 12 | 16 |
| 5' - UTR variant | 168,000 | 164,866 |
| 3' - UTR variant | 348,135 | 344,514 |
| Non-coding transcript exon variant | 277,002 | 275,909 |
| Intron variant | 12,213,614 | 12,092,056 |
| Non-coding transcript variant | 11 | 10 |
| Upstream gene variant | 878,779 | 869,207 |
| Downstream gene variant | 757,364 | 750,548 |
| Intergenic variant | 8,942,362 | 8,848,730 |

### Known trait-associated variants

The catalogue of Mendelian traits in *Sus scrofa* curated in the OMIA database (https://omia.org/home/, [22]) contained records of 47 likely causal variants (as of September 2020). However, the genomic coordinates were available for only 33 likely causal variants. Using functional annotations and sequence coverage analyses, we detected OMIA-listed variants affecting the *KIT, MC1R* and *FUT1* genes in the sequenced key ancestor animals that occurred at alternate allele frequencies between 0.013 and 1 (Supplementary Table S1, Additional File 3).

A duplication of the *KIT* gene and a splice site variant in intron 17 of the *KIT* gene are associated with the dominant white phenotype [23, 24]. Because the genotyping of larger structural and copy number variants from short-read sequencing data is notoriously difficult, we visually inspected the depth of sequencing coverage at the SSC8 region encompassing *KIT*. An increase in coverage between 41.22 and 41.78 Mb confirmed the presence of a previously reported 560 kb duplication (DUP1; Supplementary Fig. S3, Additional File 4, [25, 26]. The duplication also encompasses a copy of *KIT* that carries a splice donor site variant (SSC8: 41486012G>A, rs345599765) which manifests in a dominant white phenotype [23, 24]. The splice variant segregated at a frequency of 0.49 and 0.42 in the sire and dam line, respectively. Seven animals that carried either one or two copies of DUP1 did not carry the splice site variant and all others were heterozygous carriers. Because this variant is located within the 560 kb duplication, we observed allelic imbalance in heterozygous animals.

We detected three OMIA-listed pigmentation-associated variants in the *MC1R* gene in the sequenced pigs. All boars were homozygous carriers of a 2-bp insertion (SSC6: 182,120 - 182,121 bp), that causes a frameshift and premature translation termination, which is associated with recessive white color [27]. All animals were also homozygous carriers of two missense variants in the *MC1R* gene (SSC6: 181461T>C, ENSSSCP00000027395.1: p.Thr243Ala and SSC6: 181697A>G, ENSSSCP00000027395.1: p.Val164Ala), for which the reference alleles had been associated with red color in the Duroc breed [28].

A missense variant (SSC6: 54079560T>C; ENSSSCP00000062180.1: p.Thr102Ala; rs335979375) in the *FUT1* gene enables adhesion of enterotoxigenic Escherichia Coli F18 fimbriae (ETEC F18) to receptors at the brush border membranes of the intestinal mucosa [29]. The allele that facilitates ETEC F18 adhesion causes diarrhea in neonatal and recently weaned piglets. Since a strong selection against the ETEC F18 susceptible allele takes place in both SLW lines, we observed the disease-associated allele only in one boar from the sire line in the heterozygous state.

### Signatures of selection

We detected signatures of past selection using the composite likelihood ratio (CLR) test. Signatures of ongoing selection were identified by the integrated haplotype score (iHS) test. For both analyses, we used biallelic autosomal SNP (N_dam_ = 19,015,058, N_sire_ = 18,808,294) that were grouped into non-overlapping 100 kb windows. For the CLR tests, we considered an empirical 0.5% significance threshold to identify putative signatures of selection (Figure 3A). The number and length of candidate selection regions was higher in the dam than the sire line (14 vs. 7; 38.1 Mb vs. 26.1 Mb). Two regions on SSC3 (from 122.6 to 124.9 Mb) and SSC13 (from 140.0 to 146.1 Mb) showed evidence of selection in both lines. For the iHS analyses, we used an empirical 0.1% significance threshold to detect putative signatures of selection (Figure 3B). We detected 14 and 16 candidate regions of selection in the dam and sire line, respectively, encompassing 28.5 Mb and 32.5 Mb. Four regions on SSC1 (from 51.1 to 53.7 Mb, from 142.7 to 146.2 Mb), SSC6 (from 64.9 to 69.3 Mb) and SSC13 (from 148.0 to 150.6 Mb) were shared between both lines.

**Figure 3.**
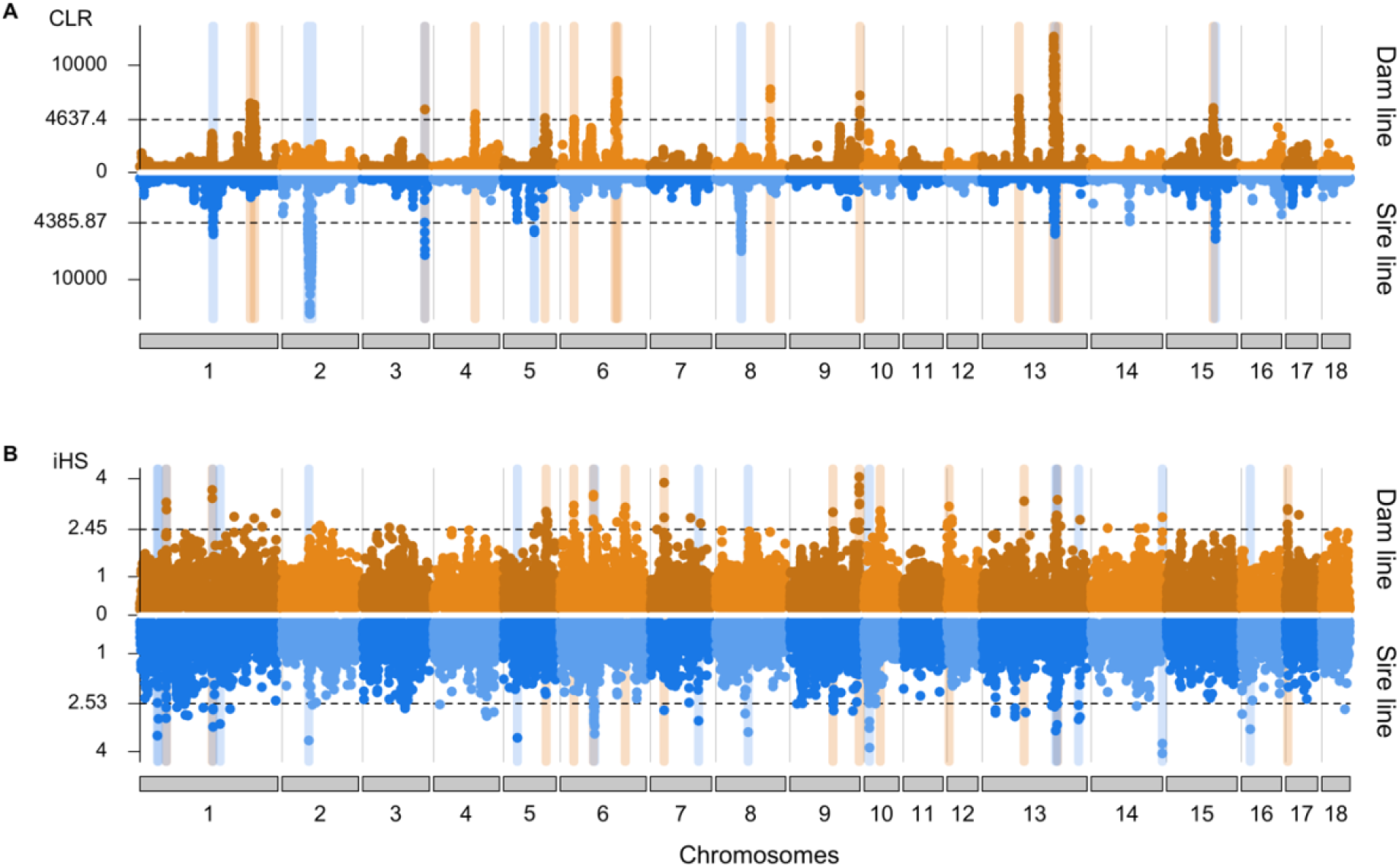
Signatures of selection detected in the sire and dam line of the SLW breed. Signatures of selection detected in the sire and dam line of the SLW breed using CLR **(A)** and iHS **(B)** Dotted lines indicate the empirical 0.5 (CLR) and 0.1% (iHS) thresholds. Blue, orange and grey vertical bars highlight signatures of selection detected in the sire, dam and both lines, respectively.

Considering both statistics, we detected more signatures of selection in the dam than sire line (28 vs. 23). Only 6 regions, detected by either CLR or iHS, overlapped between both lines. A strong signature of selection was detected in both lines with both methods on SSC13 between 140 and 152.4 Mb. The candidate region encompassed 125 genes (Supplementary Table S2, Additional File 5), as well as 63,480 and 55,835 polymorphic sites in the dam and sire line, respectively, precluding to readily prioritize candidate genes and variants responsible for the sweep.

### Reference-based genotyping from low-coverage sequencing data

In order to investigate if the 70 sequenced key ancestor animals may serve as a reference panel for genotyping by low-coverage sequencing, we sequenced the genomes of 175 pigs (84 from the sire line and 91 from the dam line) at low coverage using Gencove’s low-pass sequencing solution. The pigs also had Illumina PorcineSNP60 BeadChip-called genotypes. A principal component (Supplementary Fig. S4, Additional File 6) analysis of a genomic relationship matrix constructed from microarray-derived genotypes showed that the 175 pigs cluster with the 70 key ancestor animals.

Following quality control, we aligned a median number of 16,153,314 (between 5,950,534 and 21,168,683) read pairs (2 × 150 bp) to the porcine reference genome, achieving an average depth of coverage of 1.11-fold (from 0.38 to 1.51). On average, 54% of the reference nucleotides were covered with at least one read. Following the reference-guided low-pass sequence variant genotyping approach (GLIMPSE) proposed by Rubinacci et al. [6], we utilized the haplotypes of the 70 sequenced key ancestor animals as a reference panel to call genotypes at 22,618,811 polymorphic sites in the 175 low-pass sequenced samples.

We assessed the accuracy of genotyping by low-pass sequencing based on Illumina PorcineSNP60 BeadChip-called genotypes at 54,600 SNP, for which we were able to determine reference and alternate alleles. Of the 54,600 SNP, 6,176 and 965 were fixed for the reference and alternate allele, respectively, in the 175 pigs according to the array-called genotypes. Of 48,424 SNP that were either polymorphic or fixed for the alternate allele, 46,001 (94.99%) were also among the GLIMPSE-imputed genotypes. 2,423 SNP had microarray-derived genotypes but were missing in the GLIMPSE-imputed genotypes because these SNP were missing in the haplotype reference panel constructed from the key ancestor animals.

The genotype concordance, non-reference sensitivity and non-reference discrepancy between GLIMPSE-imputed and array-called genotypes at 46,001 autosomal SNP was 97.60, 98.73 and 3.24% in 175 low-pass sequenced pigs (Table 4, Figure 4A). When the sequence variant calling of the 175 samples was performed together with the 70 key ancestor animals using the multi-sample approach implemented in the GATK, all concordance metrics were considerably worse. Although, Beagle imputation improved the genotype calls of GATK for the low-pass sequenced samples, the genotype concordance and non-reference sensitivity was lower and non-reference discrepancy higher using GATK than GLIMPSE. Using the GLIMPSE approach improved the genotype concordance over GATK filtered & Beagle imputed variants by 13.83% and this improvement is mostly due to a lower non-reference discrepancy (Table 4).

**Table 4:** Accuracy of sequence variant genotyping in low-coverage (1.11-fold) sequencing data

| Variant genotyping approach | Genotype concordance | Non-reference sensitivity | Non-reference discrepancy |
| --- | --- | --- | --- |
| GLIMPSE | 97.60 | 98.73 | 3.24 |
| GATK raw | 75.90 | 52.35 | 30.20 |
| GATK filtered | 75.89 | 52.36 | 30.22 |
| GATK filtered & imputed | 85.74 | 96.56 | 19.34 |

**Figure 4.**
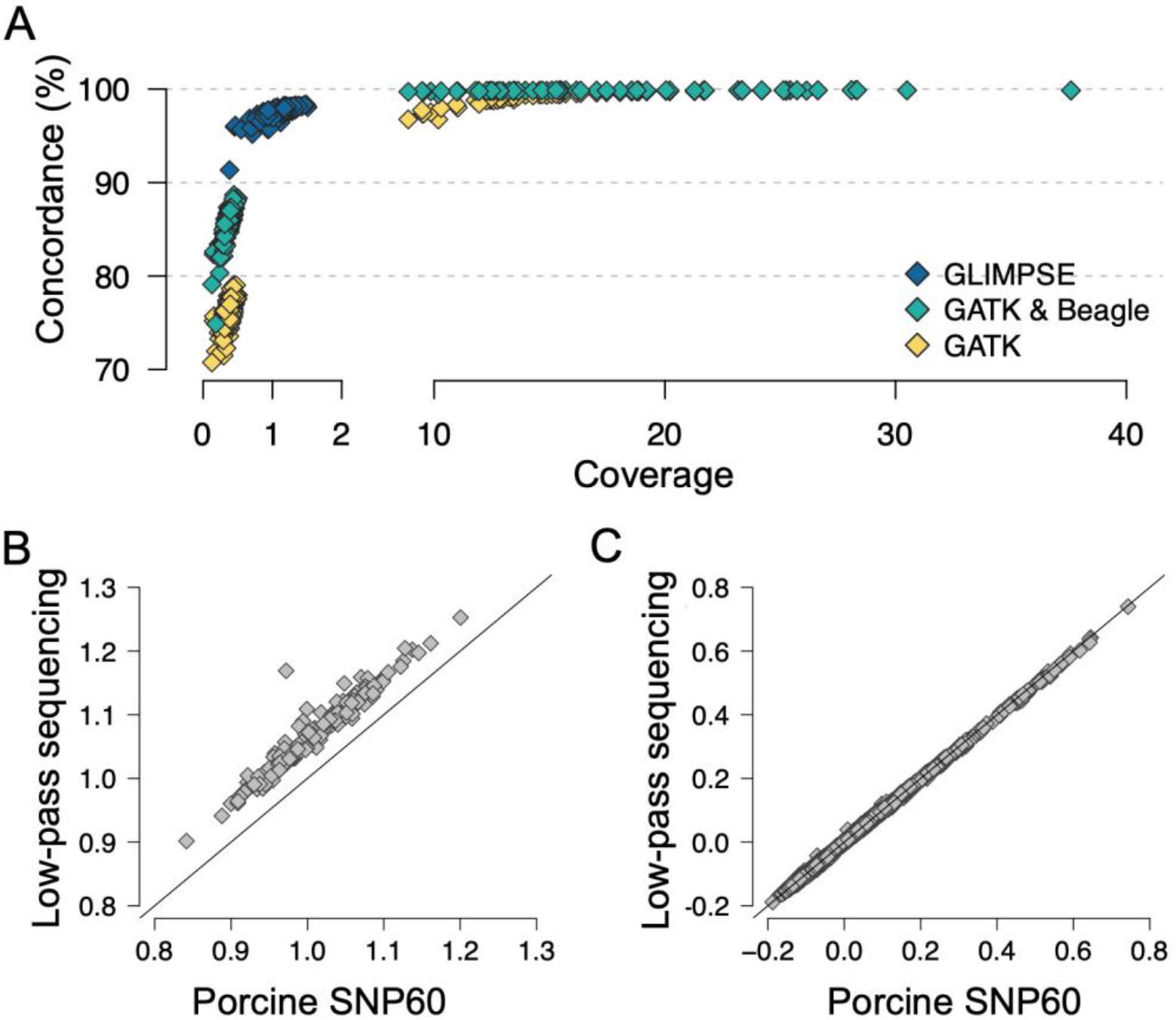
Accuracy of genotyping by low-coverage sequencing. **(A)** Concordance between array-called and sequence-called genotypes at 46,001 biallelic autosomal SNP in 243 pigs that had been sequenced at either low (N = 175; < 1.5-fold) or medium to high (N = 68; 8.88 - 37.60-fold) coverage. Correlations of (**B)** diagonal (r = 0.96) and (**C)** off-diagonal (r = 0.99) elements of genomic relationship matrices constructed from array-and GLIMPSE-called genotypes at 44,268 SNP that had minor allele frequency > 0.01.

We constructed genomic relationship matrices (GRM) from the microarray-derived and GLIMPSE-imputed genotypes of the 175 sequenced pigs based on a subset of 44,268 SNP that were detected at minor allele frequency greater than 0.01 in both datasets. Both the off-diagonal and the diagonal elements of the GRM constructed from array-derived genotypes had greater variance (σ^2^_diag_ = 3.37 × 10^−3^, σ^2^_off_ = 9.34 × 10^−3^) than corresponding elements of the GRM constructed from low-pass sequencing data (σ^2^_diag_ = 3.30 × 10^−3^, σ^2^_off_ = 9.15 × 10^−3^). While the correlation of the off-diagonal (r = 0.99) and diagonal (r = 0.96) elements was high between both GRMs, the values of the diagonal elements were higher for all samples using the GLIMPSE-imputed than microarray-derived genotypes (Figure 4B and 4C). On average, the 175 boars were homozygous for 65.58 ± 1.39% and 67.27 ± 1.49% of the 44,268 SNP when the genotypes were called from the microarray and low-pass sequencing data, respectively.

## Discussion

We applied a key ancestor animal approach to prioritize 38 and 32 boars that accounted for 95.35 and 87.95% of the genetic diversity of the SLW sire and dam line, respectively. The contributions of the SLW key ancestor animals to the current populations are considerably higher than reported for other populations. For instance, 43 key ancestor animals explained 69% of the genetic diversity of the Fleckvieh cattle population [3]. Neuditschko et al. [30] selected 41 and 55 key contributors, respectively, that explained 78% and 75% of the genetic relationship structure of the Swiss Franches-Montagnes horse and Australian Holstein-Friesian cattle population. The effective population size of the SLW sire and dam line is 44 and 72, respectively, which is less than half the effective population size of the Fleckvieh cattle and Swiss Franches-Montagnes horse population [31, 32]. Thus, a few animals that are selected based on their marginal genetic contribution to the active breeding population, account for a large fraction of the population’s haplotype diversity. It is worth mentioning that approaches other than the key ancestor animal approach may increase the haplotype diversity among the sequenced animals [33]. Nevertheless, the catalogue of 26.86 million polymorphic sites detected from the 70 sequenced boars of our study contains most alleles that segregate in the SLW populations, particularly those that occur at not too low frequency. A Ti/Tv-ratio of 2.28 indicates that the variants were of high quality [34]. In spite of the low effective population size, the nucleotide diversity (π) was high in both lines (π_dam_ = 2.24 × 10^−3^; π_sire_ = 2.23 × 10^−3^), which agrees well with estimates obtained in other European pig populations [35–37]. The nucleotide diversity in the SLW populations is higher than in cattle (1.77 × 10^−3^ – 1.90 × 10^−3^) and human (0.98 × 10^−3^ – 1.41 × 10^−3^) populations that have considerably larger current effective population sizes [38].

Although our sequencing cohort contained more animals from the sire line, we detected somewhat more autosomal variants in the dam line (N_sire_ = 23,531,919; N_dam_ = 23,774,053). While the average number of heterozygous variants detected per animal was higher in the dam line (N_sire_ = 6,180,048; N_dam_ = 6,351,565), the number of variants homozygous for the alternate allele was higher in the sire line (N_sire_ = 4,873,994; N_dam_ = 4,646,236). These differences are likely attributable to a smaller effective population size and higher genomic inbreeding in the sire line. The presence of many long ROH (> 2 Mb) suggests that recent inbreeding is higher in the sire than the dam line. Small effective population size and increasing inbreeding make both lines susceptible to the phenotypic manifestation of recessive alleles. For instance, a recessive sperm defect has recently been discovered in the sire line [39]. The management of an ever-increasing number of recessive traits is a challenge to domestic animal breeding populations [40–42]. Efficient and sustainable strategies are required to prevent the frequent manifestation of recessive diseases in populations with low effective population size.

Surprisingly, Cai et al. [43] detected fewer variants (between 20.68 and 22.11 million variants) in a considerably larger cohort of pigs (between 61 and 89) from three commercial Danish lines. Considering that Cai et al. also sequenced key ancestor animals, this difference to our study suggests higher genetic diversity in SLW. However, the depth of coverage, sequencing strategy, sequence variant genotyping and filtration approaches have major impacts on detecting polymorphic sites [44, 45]. While the effective depth of coverage realized by Cai et al. [43] is unknown to us, our samples were sequenced at an average depth of coverage greater than 16-fold. This depth of coverage enabled us to accurately detect both homozygous and heterozygous sites as evidenced by high non-reference sensitivity and genotype concordance at low non-reference discrepancy.

The principal components of a genomic relationship matrix constructed from whole-genome sequence variants revealed a separation of the animals by line. While the differentiation between the two populations might be less evident if diverse samples or an outgroup were considered in the analysis [5, 37, 46, 47], an average F_ST_ value of 0.07 corroborated that both lines diverged considerably. In fact, the average F_ST_ value observed between two SLW lines is similar to values reported between distinct European pig breeds [37, 43, 48]. The differentiation between the sire and dam line might result from distinct breeding objectives with negative genetic correlations [49]. While the sire line is mainly selected for meat and fattening traits, the dam line is mainly selected for reproduction traits. Using CLR and iHS, we detected 51 candidate signatures of selection, of which only six overlapped between both lines, suggesting that different loci are under selection in the sire and dam line. However, previous research indicates that selection for complex traits, such as production and reproduction, acts on many loci, thus barely leaves strong footprints in the genome [50, 51]. Moreover, both lines diverged only few generations ago, rendering limited time for shifts in allele frequency due to selection. We suspect that the strong differentiation between the SLW sire and dam line is also a result of genetic drift [52–54] due to very small effective population size and pronounced founder effects resulting from the unbalanced use of individual boars in artificial insemination.

A reference panel of less than 70 sequenced key ancestor animals facilitated imputing sequence variant genotypes at high accuracy and detecting trait-associated nucleotides using genome-wide association testing in cattle populations [4, 55]. Sequence variant genotypes are typically inferred using two-step imputation approaches. This requires the presence of a representative number of animals that had been genotyped at high density [9]. However, routine genotyping in the SLW populations is performed using a customized PorcineSNP60 BeadChip. Genotypes from high-density microarrays (e.g., 600K) are not available. Thus, precluding the accurate imputation of sequence variant genotypes from the key ancestor animals using the well-established stepwise imputation approach [11]. This limitation prompted us to investigate an alternative approach to reference-guided sequence variant imputation. We considered the 70 key ancestor animals as a reference to call genotypes from low-pass sequencing data (1.11-fold) of genetically similar pigs. In agreement with previous studies in human and cattle populations, the genotyping accuracy from the low-pass sequencing data was very high [6, 14, 56]. Moreover, the low-pass sequencing-derived genomic relationship coefficients were highly correlated with those obtained using microarray genotyping. This suggests that the low-pass sequencing-derived imputed genotypes may readily be used for genomic prediction [56, 57]. However, the coefficients of genomic inbreeding were higher using the genotypes from low-pass sequencing than microarray genotyping, likely because the sequenced key ancestor animals do not represent the full haplotype diversity of the SLW populations which precludes the imputation of rarer sites that predominantly occur in the heterozygous state. High-coverage sequencing of few additional animals that carry rare haplotypes may mitigate this ascertainment bias [58] and increase the accuracy of genotyping by low-pass sequencing, particularly for rare alleles. While a subset of the 22.62 million variants obtained is sufficient to accurately predict genomic breeding values, the full variant catalogue, once available for a large mapping cohort, will facilitate powerful genome-wide association studies at nucleotide resolution.

## Conclusions

The high-coverage sequencing of 70 key ancestor animals from two SLW lines and subsequent reference-guided variant discovery revealed 26,862,369 polymorphic sites. Population-genetic analyses suggest considerable genetic differentiation between both lines. Our results indicate that the key ancestor genomes may serve as a haplotype reference panel for genotyping by low-pass sequencing at high accuracy in the Swiss pig breeds. Using genotyping by low-pass sequencing increases the variant density over the currently used microarray by > 350-fold, thus providing an valuable resource for powerful genome-wide association testing.

## Methods

### Animals and whole-genome sequencing

Whole genome sequence data were generated for 70 boars. Sixty-five boars (32 from the dam line and 33 from the sire line) were selected based on their marginal genetic contribution to the current breeding populations using a key ancestor approach [2]. The marginal genetic contribution was estimated based on a numerator relationship matrix that was constructed using the PyPedal python package [59]. The effective population size of the sire and dam line was estimated based on the difference in pedigree-derived inbreeding coefficients between active breeding animals and their parents that were extracted from the numerator relationship matrix. Animals born after 01.01.2018 were considered as active breeding animals. In addition, we considered whole-genome sequence data from five boars from the sire line that were generated previously [39]. DNA was prepared from preserved blood samples that were provided by SUISAG (the Swiss competence center for pig breeding). No animals were specially sampled for the present study. Illumina TruSeq PCR-free libraries with insert sizes of 350 bp were prepared and sequenced with an Illumina NovaSeq6000 instrument using 2×150 bp paired-end reads.

### Alignment quality, read mapping and depth of coverage

We used the fastp software [60] to remove adapter sequences and reads that had Phred-scaled quality less than 15 for more than 15% of the bases. Subsequently, the filtered reads were aligned to the SSC11.1 assembly of the porcine genome [61] using the mem-algorithm of the BWA software [62]. The Picard tools software suite [63] and Sambamba [64] were applied to mark duplicate reads and sort the alignments by coordinates, respectively. To calculate depth of coverage, we extracted the number of reads covering a genomic position using the mosdepth software [65]. For the coverage calculation, we discarded reads with mapping quality < 10 and SAM bitwise flag value of 1796.

### Variant calling

We used the BaseRecalibrator module of the Genome Analysis Toolkit (GATK - version 4.1.0; [17]) to adjust the base quality scores while supplying 63,881,592 unique positions from the porcine dbSNP version 150 as known variants. We applied the HaplotypeCaller, GenomicsDBImport and GenotypeGVCFs modules from the GATK to discover and genotype SNP and INDEL in the 70 SLW pigs together with 28 samples from various breeds that were sequenced earlier. Subsequently, we applied the VariantFiltration module of the GATK according to best practice recommendations for site-level hard filtration to retain high-quality variants. Beagle (version 4.1; [16]) haplotype phasing and imputation was applied to impute sporadically missing sites and improve the primary genotypes obtained using the GATK.

The concordance between sequence-and array called genotypes was calculated for 68 pigs that also had Illumina PorcineSNP60 BeadChip microarray-derived genotypes. We considered only autosomal SNP. We converted the TOP/BOT alleles of the microarray-derived genotypes to REF/ALT allele coding to make them compatible with the sequence-derived genotypes. This was possible for 54,600 SNP. Sequence variant genotyping accuracy was quantified using genotypic concordance, non-reference sensitivity and non-reference discrepancy [17, 44].

Invariant sites and variants within regions with an excessive depth of coverage (> mean coverage + 2 * SD) were removed using VCFtools (v. 0.1.16; [66]). The resulting data were split into two datasets containing 23,774,053 and 23,531,919 variants segregating in 32 boars from the dam line and 38 boars from the sire line, respectively.

### Functional annotation

Functional consequences of the variants (including SIFT scores [21] for missense variants) were predicted with the Ensembl Variant Effect Predictor (VEP, version 91.3; [20]) using local cache files from Ensembl release 98. The transition to transversion ratio (Ti/Tv) was calculated using BCFtools command *stats* (version 1.8; [67]).

### Detection of mendelian trait-associated variants and coverage analysis

We downloaded genomic coordinates of 47 likely causal variants from the Online Mendelian Inheritance in Animals (OMIA) database [22]. Genes harboring likely causal variants for which the genomic coordinates were not annotated according to SSC11.1 were manually inspected. Read alignments and sequence coverage in regions harboring known larger structural variants were manually inspected.

### Population structure and genetic diversity analysis

The structure of the two lines was investigated using ADMIXTURE (v1.3.0; [68]). To avoid confounding due to extensive linkage disequilibrium (LD), we removed correlated loci based on high levels (r^2^ > 0.6) of pairwise LD using PLINK (version 1.9; [69]) with the “--indep-pairwise 100 25 0.6” option before running the ADMIXTURE analysis. The number of ancestral clusters (K) was set from 1 to 3, and five-fold cross-validation was performed to determine the K value with the lowest cross-validation error.

A genomic relationship matrix was built using 23,691,198 autosomal sequence variants that had a minor allele frequency higher than 0.01 using PLINK. The principal components of the genomic relationship matrix were calculated using the GCTA (version 1.92.1; [70]) software. We applied the GCTA flag “--grm-singleton” to identify four pairs of animals with relationship coefficients ranging from 0.32 to 0.37. One animal from each pair was removed for the F_ST_ and signature of selection analyses (1 from the dam line and 3 from the sire line).

We calculated the weighted genome wide fixation index (F_ST_, [18]) based on pairwise differences in the variances of allele frequencies using 24,926,366 biallelic variants. F_ST_ values were calculated in 10 kb sliding windows with an overlap of 5 kb using the “--weir-fst-pop” flag of VCFtools (v.1.2.11; [66]). The manhattan plot was constructed using the R package qqman [71].

Nucleotide diversity (π) was calculated over all biallelic autosomal variants in 10 kb sliding windows with an overlap of 5 kb using VCFtools.

Runs of homozygosity (ROH) were estimated with BCFtools/ROH [19] using the GATK-derived genotypes (containing the Phred-scaled likelihoods). We considered biallelic SNP that had non-missing genotypes in all animals (maximal missing count per site was set to 0). According to Tortereau et al. [72], we assumed a constant recombination rate of 0.7 cM/Mb along the chromosomes. Average genomic inbreeding (F_ROH_) was calculated assuming an autosomal genome length of 2,265,774,640 bases. Following a recent study by Bhati et al. [73], we classified the ROH based on their length (short: 50 - 100 kb, medium: 100 kb - 2 Mb, long: > 2 Mb).

### Signatures of selection (CLR and iHS) and candidate regions

Putative signatures of selection were detected using integrated Haplotype Scores (iHS) and composite likelihood ratios (CLR). The iHS [74] reveals ‘soft sweeps’, i.e., signatures of selection where selection for beneficial alleles is still ongoing. The CLR [75] reveals ‘hard sweeps’, i.e., signatures of selection where beneficial alleles recently reached fixation. We considered 24,926,366 autosomal biallelic SNP from 31 and 35 boars from the dam and sire line, respectively. The genotypes were phased using Beagle (version 5.1; [76]) with disabled imputation and effective population size set to 50. The CLR statistic was calculated chromosome-wise with the SweepFinder2 software [77] using a pre-computed empirical allele frequency spectrum and 100 kb spacing between test sites (-lg 100000). Using the R package rehh 2.0 [78], we applied the function *scan_hh* to estimate the integrated extended haplotype homozygosity (EHH) on variants with MAF > 0.05 for each chromosome separately. Subsequently, we applied the function *ihh2ihs* to obtain standardized iHS values in 100 kb non-overlapping windows.

The function *calc_candidate_regions* from the rehh 2.0 package [78] was applied to select candidate signatures of selection in 100 kb windows using the parameters “window_size = 1E6”, “overlap = 1E5”, “pval = F” and “min_n_extr_mrk = 1”. Empirical significance thresholds were chosen after visual inspection of the distribution of the test statistics (0.1% in iHS and 0.5% in CLR). Genes overlapping with candidate signatures of selections were determined based on the Ensembl (release 98) annotation of the porcine genome.

### Analysis of low-pass sequence data

A median number of 16,131,419 paired-end (2×150bp) reads were generated for 96 pigs from the dam line and 96 pigs from the sire line. Adapter sequences and bases and reads with low sequencing quality were removed with fastp [60]. Subsequently, the reads were aligned to the porcine reference genome (SSC11.1) using the mem-algorithm of BWA [62] and duplicate reads were marked using Samblaster [79]. Following the read alignment, six samples were excluded because the mapping rate and the proportion of properly paired reads was less than 70 and 75%, respectively. Additionally, we excluded 10 samples for which the average coverage was less than 0.2-fold and one sample for which ancestry could not be verified.

To compile the reference haplotypes, we retained 22,618,811 biallelic autosomal SNP that were polymorphic (minor allele count ≥ 1) among the 70 key ancestor pigs. Following the approach proposed by Rubinacci et al. [6], we used the *mpileup* and *call* commands of BCFtools [67] to calculate genotype likelihoods at the 22,618,811 polymorphic sites in the 175 low-pass sequenced and reference-aligned samples. Subsequently, we applied the phasing and imputation algorithm implemented in GLIMPSE_phase [6] to refine the BCFtools-derived genotype calls using the previously established haplotype reference panel. This approach produced genotypes at 22,618,811 sites for the 175 low-pass sequenced samples. A genomic relationship matrix among the low-pass sequenced animals was constructed from the low-pass sequencing data-derived genotypes using GCTA [70].

## List of abbreviations

CLR: Composite likelihood ratio
iHS: Integrated haplotype score
INDEL: Insertions and deletions
LD: Linkage disequilibrium
MAF: Minor allele frequency
OMIA: Online Mendelian Inheritance in Animals
PCA: Principal component analysis
QTL: Quantitative trait loci
ROH: Run of homozygosity
SLW: Swiss Large White
SNP: Single nucleotide polymorphism
Ti/Tv: Transition/transversion ratio
WGS: Whole-genome sequencing

## Supplementary information

Additional File 7: Additional files

PDF – Information of Additional files 1 to 6

## Declarations

### Ethics approval and consent to participate

No animals were sampled for the present study. Thus, no ethical approval was required.

### Consent for publication

Not applicable.

### Availability of data and materials

Raw sequencing read data of all key ancestor animals are available at the European Nucleotide Archive (ENA) (http://www.ebi.ac.uk/ena) of the EMBL at BioProject PRJEB38156 and PRJEB39374.

### Competing interests

AH is employee of SUISAG (the Swiss pig breeding and competence centre). All other authors declare that they have no competing interests.

## Funding

This study was financially supported by SUISAG, Micarna SA and the ETH Zürich Foundation. The funding body was not involved in the design of the study and collection, analysis, and interpretation of data and in writing the manuscript.

## Authors’ contributions

Analysis of whole-genome sequencing data: AN MB NKK DC HP; Conceived and designed the experiments: AN HP; Conceptualisation: HP AH; Secured funding: HP SN; Wrote the paper: AN HP; Critically revised the manuscript: all authors; Read and approved the final version of the manuscript: all authors.

## Acknowledgements

We acknowledge Gencove (https://gencove.com/) for providing support to obtain low-pass sequencing data.

