## Supplementary figures for "Characterization of a haplotype-reference panel for genotyping by low-pass sequencing in Swiss Large White pigs"

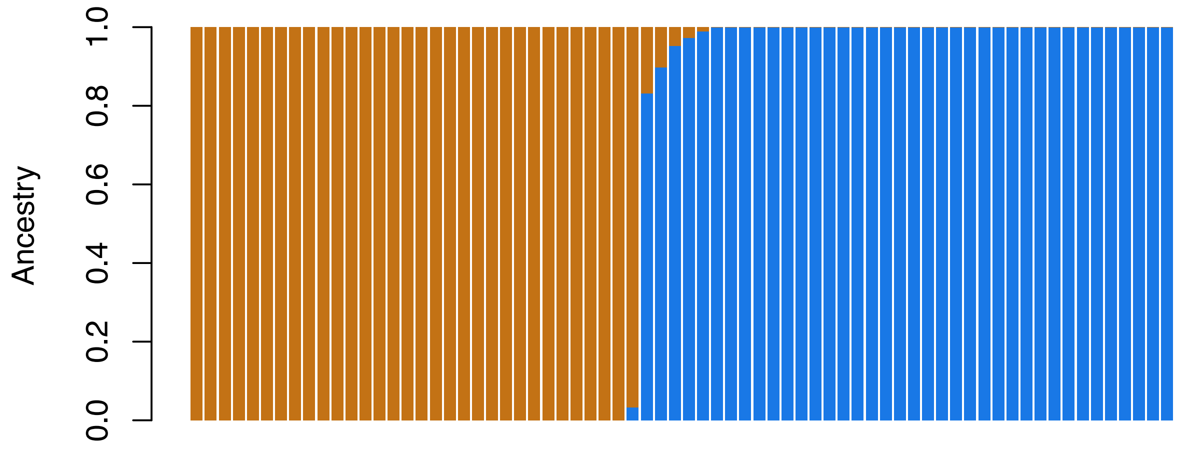


**Additional File 1: Fig. S1:** Admixture analysis.

Ancestry of 70 pigs with K = 2 ancestral populations estimated using 1,207,189 biallelic SNPs after LD pruning. Ancestry proportions were estimated using the ADMIXTURE software. Each bar represents an individual and the colors indicate the proportion of genes originating from K ancestral populations. The animals are ordered by population.

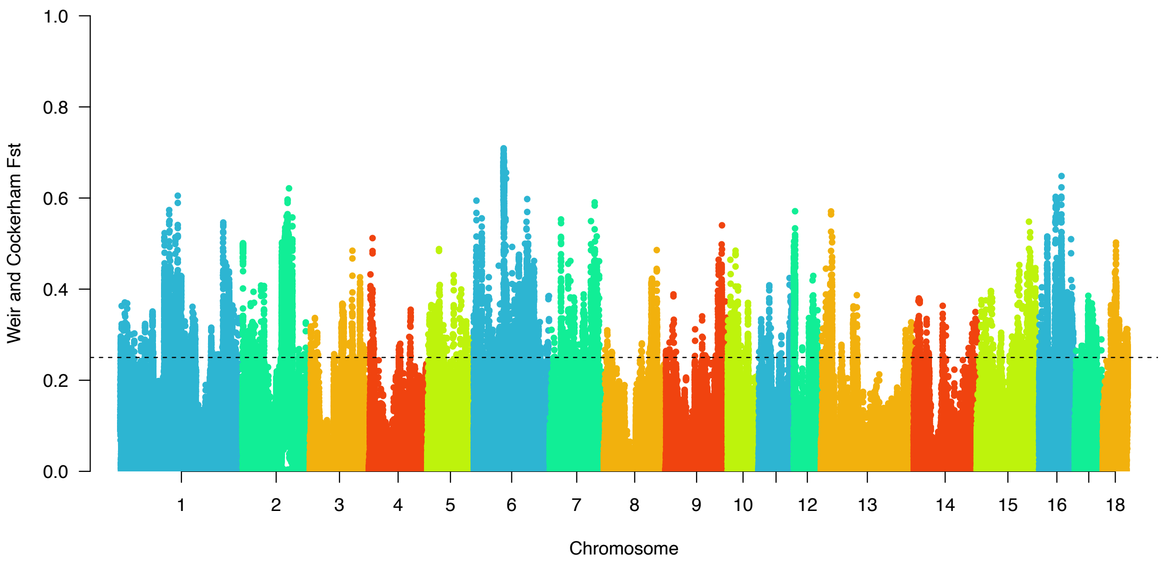


**Additional File 2: Fig. S2:** Manhattan plot of $F_{ST}$ values.

Weir and Cockerham $F_{ST}$ estimates were calculated in 10 kb sliding windows between 31 dam and 35 sire boars. Black dotted line indicates a value of 0.25.

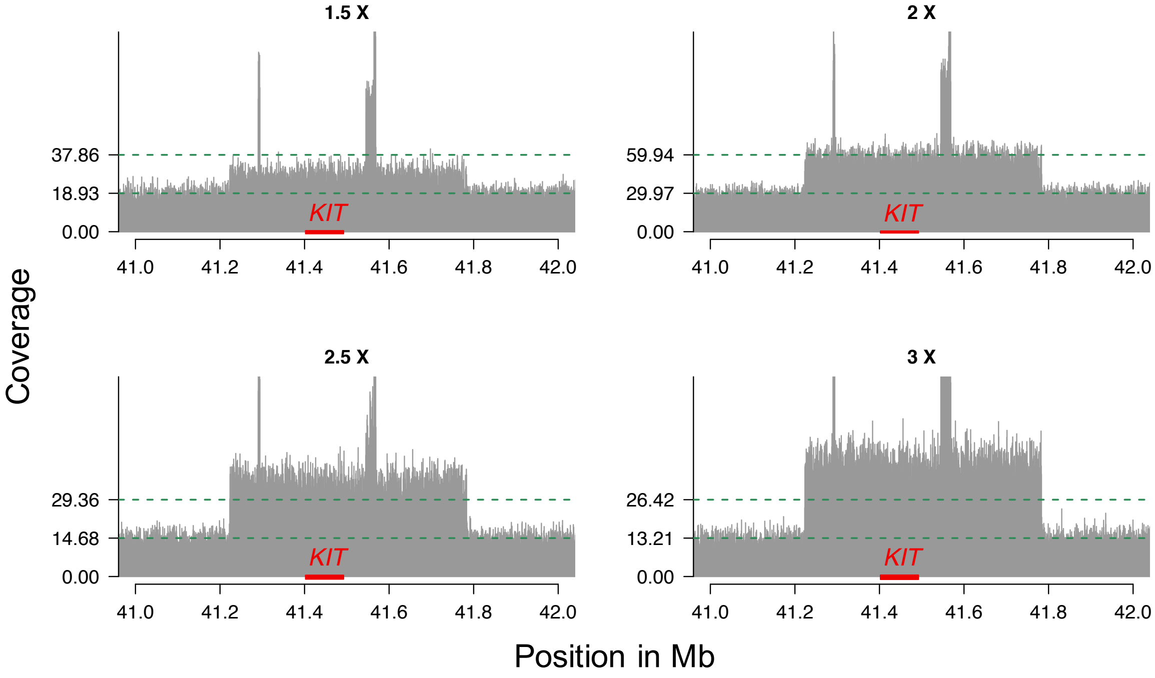


**Additional File 4: Fig. S3:** Depth of coverage at a region on SSC8 encompassing the *KIT* gene.

Representative plots of different depth of coverage detected in the sequenced pigs at a large duplication (DUP1 - SSC8: 41,223,212 - 41,783,660 bp) encompassing the *KIT* gene. Grey vertical bars represent the absolute coverage observed in four animals. The green dotted lines represent the median and 2*median coverage along SSC8. In order to determine the number of extra copies, we divided for each sequenced animal the average coverage observed at SSC8 by the average coverage observed at DUP1 (chr8: 41,223,212 - 41,783,660 bp). The number of additional copies ranged from 1 to 4. 16, 27, 4, 20, 2 and 1 animal had 1.5 - 1.9x, 2x, 2.1 - 2.4x, 2.5 - 2.9x and 3x the average coverage of SSC8 at DUP1, respectively. The average copy number was 2.07 and 2.19 in the dam and sire line, respectively. The 560 kb duplication (DUP1) encompasses two smaller duplications DUP2 and DUP3/4. DUP2 is 4.3kb long and upstream, while DUP3 and DUP4 are 23kb and 4.3kb duplications downstream of the *KIT* gene.

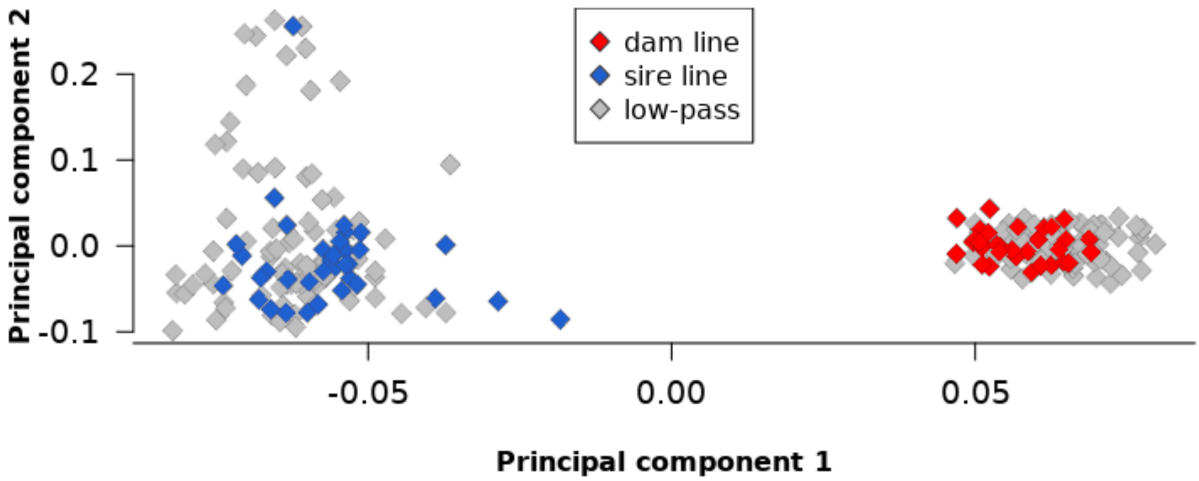


**Additional File 6: Fig. S4:** Principal components analysis of key ancestor and low-pass sequenced animals.

Plot of the first two principal components showing the relationship of 96 dam and 96 sire animals sequenced at low (< 1.5-fold) coverage and 32 dam and 38 sire animals sequenced at high (~ 16.5-fold) coverage.

### Supplementary tables

**Additional File 3: Table S1:** List of variants listed in the OMIA database and their corresponding frequency in the two pig lines.

**Additional File 5: Table S2:** Candidate signatures of selection based on CLR and iHS analyses.

Genes annotated to the region are given for each signature of selection.
